# A new pathway of pre-tRNA surveillance in yeast

**DOI:** 10.1101/561944

**Authors:** Matthew J. Payea, Thareendra C. De Zoysa, Eric M. Phizicky

## Abstract

During tRNA maturation in yeast, aberrant pre-tRNAs are targeted for 3’-5’ degradation by the nuclear surveillance pathway, and mature tRNAs are targeted for 5’-3’ degradation by the rapid tRNA decay (RTD) pathway, due to lack of certain body modifications or to destabilizing mutations. Here we show that the RTD pathway also targets pre-tRNAs through an unknown, but distinct, mechanism that occurs after nuclear export. Anticodon stem RTD variants of both tRNA^Tyr^ and tRNA^Ser(CGA)^ are substrates for pre-tRNA RTD, triggered by the accumulation of end-matured unspliced pre-tRNA due to altered secondary structure of the region comprising the anticodon stem-loop and the intron. Furthermore, increased nuclear availability of a pre-tRNA RTD substrate can provoke decay by nuclear surveillance. We interpret these results in terms of a model of opportunistic tRNA decay, wherein tRNAs are degraded due to a combination of structural instability and increased availability to decay pathways.

## INTRODUCTION

In all organisms, the central role of tRNAs in decoding mRNA during translation places restrictions on the properties of tRNA. tRNAs must adopt a conserved structure to uniformly participate in translation (1–3); they must be selective in charging by their cognate synthetase (4–6), and in their anticodon-codon interactions (7–11); they must be flexible enough for accommodation and passage through the ribosome (12–16); and they must be stable enough to have long half-lives, enabling participation in multiple rounds of translation (17–20).

tRNAs undergo a highly complex maturation process. In the yeast *Saccharomyces cerevisiae* and other eukaryotes, tRNAs are transcribed by Pol III as pre-tRNAs with a 5’ leader, a 3’ trailer, and in some cases an intron, all of which must be removed before the tRNA can participate in translation (21). In vertebrates and higher eukaryotes, these end trimming and splicing steps are completed in the nucleus (22,23), but in *S. cerevisiae* and *Schizosaccharomyces pombe*, intron-containing pre-tRNAs are exported from the nucleus after end trimming and then spliced in the cytoplasm on the surface of the mitochondria by the heterotetrameric splicing endonuclease (SEN) complex (24–26). In addition, an extensive set of modifications are added to the body and anticodon loop throughout the course of processing (22,27,28), including a final set of modifications that occur after splicing, after retrograde import of spliced tRNAs into the nucleus, or after re-export of the tRNAs into the cytoplasm (29–32).

The necessity for proper tRNA maturation and function is evident from the numerous human diseases associated with incorrectly processed tRNAs. Hypomodification of tRNAs, caused by mutation of either the modifying enzyme or the tRNA itself, can result in numerous mitochondrial and neurological diseases (33–40); defects in tRNA splicing caused by mutations in the human SEN complex can cause pontocerebellar hypoplasia and neurodegeneration (41,42); and mutations in the aminoacyl tRNA synthetases can cause a host of neurological diseases (43,44).

In *S. cerevisiae*, the fidelity of tRNA maturation is monitored both as pre-tRNA, by the nuclear surveillance pathway, and as mature tRNA, by the rapid tRNA decay (RTD) pathway. The nuclear surveillance pathway acts through the TRAMP complex and the nuclear exosome to degrade pre-tRNA_i_^Met^ lacking the m^1^A modification (18,45–47), and may also degrade as much as half of the total transcribed pre-tRNA, presumably due to random mutations or folding defects (17). After maturation, tRNAs are subject to another round of surveillance by the rapid tRNA decay (RTD) pathway, which degrades certain hypomodified tRNAs and tRNAs with destabilizing mutations in the acceptor/T-stem, employing the 5’-3’ exonucleases Rat1 in the nucleus and Xrn1 in the cytoplasm (19,20,48–50). The RTD pathway targets tRNAs due to exposure of the 5’ end arising from reduced stability of the acceptor/T-stem, as shown by both structure probing and Xrn1 assay of hypomodified tRNA^Ser(CGA)^ (tS(CGA)) and tS(CGA) variants, and by Xrn1 assay of hypomodified tV(AAC) (48). The degradation of all identified substrates of the RTD pathway is suppressed by a *met22Δ* mutation, likely due to elevated levels of the metabolite 3’, 5’ adenosine bisphosphate (pAp) found in a *met22Δ* strain, which inhibits both Xrn1 and Rat1 *in vitro* and *in vivo* (20,51,52). There is also evidence that the RTD pathway may be conserved in humans, since Xrn1 and the human Rat1 homolog Xrn2 can mediate the degradation of tRNA_i_^Met^ after heat shock (53).

Although the RTD pathway targets substrates due to 5’ end exposure (48), it was surprising to find, from high throughput analysis of variants of the suppressor *SUP4_oc_* (tRNA^Tyr(GUA)^ with a UUA anticodon), that the RTD pathway could also target variants with destabilizing mutations in the anticodon stem. Such anticodon stem mutations were not expected to significantly destabilize the 5’ end and trigger RTD (54,55).

Here we show that RTD of anticodon stem variants is quantitatively equivalent to RTD of acceptor stem variants *in vivo*, despite the much larger exposure of acceptor stem variants to 5’-3’ exonucleases *in vitro*. We explain this discrepancy by showing that anticodon stem RTD substrates have a defect in pre-tRNA maturation, causing accumulation of unspliced pre-tRNA that is degraded by the RTD pathway through an unknown mechanism, as it is not inhibited by deletion of *XRN1* or mutation of *RAT1*. We provide evidence that mutations that restore pre-tRNA structure reduce pre-tRNA accumulation and pre-tRNA RTD. Furthermore, we find that pre-tRNA RTD and anticodon stem RTD is general since it also occurs in both tY(GUA) and tS(CGA) variants. We explain these results in terms of a model of opportunistic decay in which destabilizing variants are subject to surveillance at each step of tRNA biogenesis and after maturation, and are degraded due to a combination of instability and availability of each tRNA intermediate.

## MATERIALS AND METHODS

### Yeast strains

The yeast strains used are shown in Supplementary Table 1. Deletions were introduced by standard PCR amplification of the appropriate strain from the YKO collection (56) (or of a derivative strain with a different drug marker), followed by linear transformation, and PCR of transformants to confirmation the deletion. The *rat1-107* allele was introduced with a *URA3* marker as described previously (20).

### Construction and integration of variants of *SUP4_oc_, tS(CGA), tV(AAC/CGA)*

*SUP4_oc_* variants were constructed by annealing of overlapping oligomers (IDT) followed by ligation into the Bgl II Xho I site of the tRNA gene cassette of plasmid AB230-1, as previously described (54). tS(CGA) variants and tV(AAC/CGA) were constructed in a similar manner. tRNA variants were integrated at the *ADE2* site of strains by linear transformation of a Stu I fragment containing the tRNA cassette, the adjacent *HIS3* marker, and flanking *ADE2* sequences, deleting part of *ADE2* (48). Three biological isolates of each strain were isolated and used for all of the experiments reported, unless otherwise noted.

### Growth of cells for isolation of bulk RNA

Three independent isolates of strains with integrated tRNA variants were grown overnight at 28°C in 5 mL YPD media (1% yeast extract, 2% peptone, 2% dextrose, supplemented with 80 mg/l adenine hemisulfate), inoculated into 5 ml fresh media at OD_600_ ~ 0.1, and grown to an OD ~1, and then 2 OD samples were harvested by microfugation, washed with 1 mL water, frozen on dry ice, and stored at −80°C. Then bulk RNA was extracted using hot phenol as described previously (57), followed by addition of 20 μg glycogen, phenol-chloroform-isoamyl alcohol (PCA) extraction, two ethanol precipitations, and resuspension in ddH_2_O.

### Purification of Xrn1 and the Rat1/Rai1 complex

Xrn1 was purified from yeast strain BCY123 (58), after galactose-induced expression of a P*_GAL1_-XRN1* construct in which *XRN1* was fused at its 3’ terminus to the PT tag, bearing a protease 3C site-HA epitope-His6-ZZ domain of protein A, as described previously **(48)**. Rat1/Rai1 was purified analogously from a dual *P_GAL1,10_* expression plasmid expressing *RAT1-PT* and His10-RAI1 **(59)**.

### Purification of tRNAs

*SUP4_oc_* variants were purified together with tRNA^Tyr^ using 1.0 mg bulk RNA from the corresponding *met22Δ* strain and the biotinylated oligomer MP 129 (nt 76-53), as previously described (48,60).

### *In vitro* digestion with exonucleases

Purified tRNAs were analyzed for exonuclease sensitivity essentially as previously described (48). A purified mixture of tRNA^Tyr^ and *SUP4_oc_* (7.5 ng) was melted at 95 °C for 10 minutes in buffer containing 25 mM Tris pH 8.0 and 150 mM NaCl, immediately placed on ice, refolded at 37°C for 20 minutes in buffer containing 13.9 mM Tris pH 8.0, 83.3 mM NaCl, 2.2 mM MgCl_2_, 0.55 mM DTT, 0.11 mg/mL BSA, and then incubated as indicated with purified Xrn1 or Rat1/Rai1 in buffer containing 4.5 mM Tris pH 8.0, 90 mM NaCl, 2.1 mM MgCl_2_, 0.6 mM DTT, 0.1 mg/mL BSA, 5% glycerol. Reactions were stopped on dry ice, extracted with PCA, ethanol precipitated with 20 μg of glycogen, and then resuspended in ddH_2_O.

### Poison primer extension analysis of *SUP4_oc_* variants

To analyze relative amounts of *SUP4_oc_* and tRNA^Tyr^ in bulk RNA or purified mixtures, we used a poison primer extension assay with an appropriate primer (Supplementary Table 2) and ddCTP or ddTTP, essentially as previously described (54). Briefly, ~0.5 pmol of 5’ radiolabeled primer and ~1.0 μg of bulk RNA was denatured by heating at 95 °C and then annealed in water, followed by primer extension with 2U AMV reverse transcriptase (Promega) in supplied buffer containing 1 mM ddNTP, and 1 mM remaining dNTPs, incubation for 1 hour at 50°C, and transfer to −20°C. Aliquots of reactions were diluted 2-fold in formamide dye, heated at 95°C for 3 minutes, and resolved on a 15% polyacrylamide gel (29:1) containing 7 M urea in 1 X TBE. The gel was dried on a Model 583 Biorad gel dryer at 78°C for 40 minutes, exposed and analyzed using an Amersham Typhoon phosphoimager and Image Quant v5.2.

The relative amount of *SUP4_oc_* variant in each sample is expressed as % tY(GUA) (% tY), calculated as the quotient of the signal for the *SUP4_oc_* variant (typically the G_30_ stop) divided by the endogenous tRNA^Tyr^ signal (typically the G_34_ stop), each first corrected for background. The % tY was averaged over three biologically independent strains grown and analyzed in parallel, and the error calculated as a standard deviation. The RTD ratios were calculated as the quotient of the average % tY value in a *met22Δ* strain divided by the corresponding average % tY value in a *MET22*^+^ strain (analyzed in the same gel from primer extensions done at the same time), with the error propagated appropriately for a division operation.

We note that despite some variability in % tY from experiment to experiment, the RTD ratios obtained from *MET22*^+^ and *met22Δ* strains grown in triplicate and analyzed at the same time are largely consistent between experiments. The variability in % tY likely arises from variations in the hybridization efficiencies of the primer extension probe to *SUP4_oc_* variants in different sets of assays. This results in variable % tY measurements for the same bulk RNA preparations assayed at different times, but the same % tY from biological triplicates assayed at the same time.

### Northern analysis of *SUP4_oc_* Variants

For northern analysis, independently made strains were grown in parallel, bulk RNA was isolated, and for each sample, 1 μg RNA was mixed with an equal volume formamide dye, heated at 60°C for 5 minutes, and resolved on a 10% polyacrylamide (19:1), 7M urea 1X TBE gel, which was transferred onto a Amersham Hybond−-N^+^ membrane (GE Healthcare), crosslinked on a UV Stratalinker 2400, and hybridized to probes (~ 12 pmol 5’ ^32^P-labeled DNA oligomer) as indicated. Oligomers are described in Supplementary Table 3.

### Acidic Northern analysis of *SUP4_oc_* Variants

To analyze aminoacylated tRNAs, RNA was prepared from cells with glass beads under acidic conditions at pH 4.5, and Northern analysis was done as described previously (19). RNA was resolved on a 6.5% polyacrylamide (19:1), 8 M urea, 0.1 M sodium acetate (pH 5.0) gel and separated at 4°C for approximately 12 hours at 600 volts. The gel was transferred in 0.1M sodium acetate buffer onto an Amersham Hybond−-N^+^ membrane followed by crosslinking and hybridization as described above.

## RESULTS

### Anticodon stem variants are susceptible to RTD *in vivo* but not to RTD exonucleases *in vitro*

Because of our finding that numerous anticodon stem variants could provoke RTD in the absence of known mechanisms for them to significantly destabilize the tRNA 5’ end (54), we quantitatively compared RTD of anticodon stem and acceptor stem *SUP4_oc_* variant substrates. We examined the tRNA levels of biological triplicates of three acceptor stem RTD variants, seven anticodon stem variants, and one anticodon loop variant (Figure 1A,B, and C). We used poison primer extension to directly compare tRNA levels of *SUP4_oc_* variants as a percentage of endogenous tY(GUA) in a *MET22*^+^ strain and in a *met22Δ* strain, in which the 5’-3’ exonucleases of the RTD pathway are inhibited, and then calculated the ratio of these two values (*met22Δ/MET22*^+^) to obtain the RTD ratio (54,61). Consistent with previous findings, we found that a *met22Δ* mutation resulted in a large quantitative increase in the tRNA levels for all 11 tested variants and thus a substantial RTD ratio (Figure 1C and D). Seven variants had an RTD ratio greater than 4 (C_5_U, A_29_U, G_30_A, C_40_U, U_41_A, U_42_A, G_69_C), and four had an RTD ratio between 2 and 4 (U_2_C, A_28_U, A_31_U, C_32_A), indicating that they were all RTD substrates. Moreover, the average RTD ratio of anticodon stem variants (5.4, or 5.0 counting the C_32_A anticodon loop variant) was similar to, or slightly greater than, that of the acceptor stem variants (4.4). This finding emphasizes the unusual nature of anticodon stem RTD substrates in that despite their predicted inability to substantially increase exposure of the 5’ end to exonucleases, they provoke RTD *in vivo* at least as effectively as mutations in the acceptor stem. The C_32_A variant is particularly notable as an RTD substrate since the C32 residue is in the anticodon loop immediately adjacent to the anticodon stem and, if anything, is predicted to slightly improve stability of the anticodon stem-loop (62).

**Figure 1.**
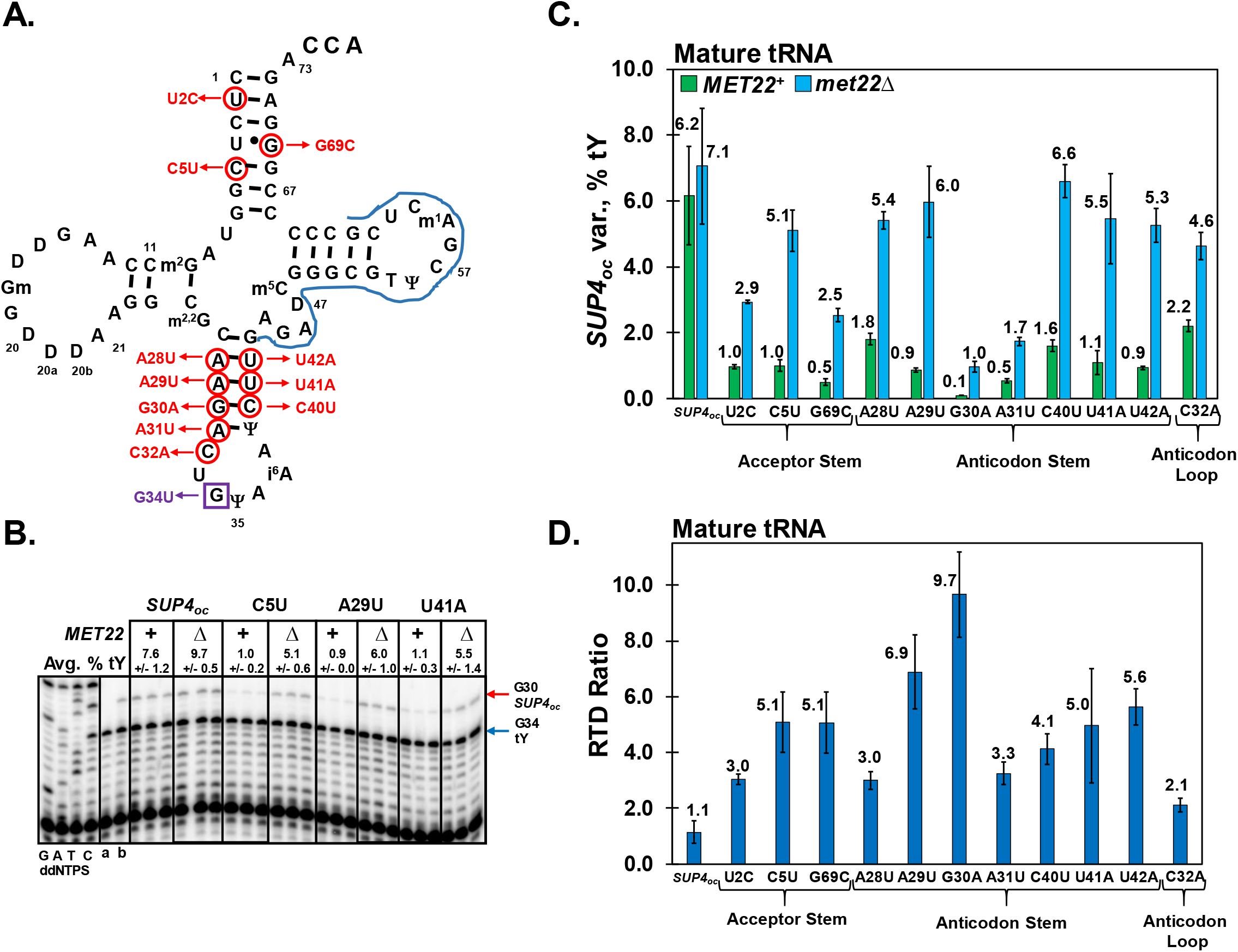
The RTD of anticodon stem *SUP4_oc_* variants is as quantitatively significant as that of acceptor stem variants. **(A) Schematic of *SUP4_oc_* secondary structure and variants examined.** The secondary structure of tY(GUA) is shown, together with the G_34_U ochre mutation, purple box; variants examined, red circles; and P7 primer (complementary to nt 62-43) used for poison primer extension analysis of mature tRNA in (B), blue line. **(B) Poison primer extension analysis of *SUP4_oc_* anticodon stem and acceptor stem variants.** Strains as indicated containing an integrated copy of a *SUP4_oc_* tRNA were grown at 28°C to mid-log phase, and bulk RNA was analyzed by poison primer extension with the P7 primer (62–43) in the presence of ddCTP, producing a stop at G_34_ for endogenous tY(GUA) (blue arrow) and at G_30_ for integrated *SUP4_oc_* variants (red arrow). Samples were resolved on a 15% polyacrylamide 7M urea gel. A sequencing ladder is shown at the left. To confirm that all the measured G_30_ signal is from *SUP4_oc_* variants, bulk RNA was also analyzed from a strain containing no *SUP4_oc_* variant, lane a; and an intronless *SUP4_oc_* variant, lane b. *SUP4_oc_* variants were quantified as described in Materials and Methods as a percentage of the endogenous tY(GUA) (% tY) and an average value was determined for each *SUP4_oc_* variant from the triplicate samples in each genetic background (Avg. % tY). As noted in more detail in Materials and Methods, data compared for primer extensions done at the same time, are highly consistent. **(C) *SUP4_oc_* anticodon stem and acceptor stem variants have increased tRNA levels in *met22Δ* relative to *MET22*^+^.** Bar graph depicting average % tY values with associated standard deviations for variants analyzed (For *SUP4_oc_*, n = 12 across four different experiments each with three biological variants, for all other variants n = 3); *MET22^+^*, green, *met22Δ*, blue. **(D) *SUP4_oc_* anticodon stem and acceptor stem variants are similarly degraded by the RTD pathway *in vivo*.** Bar graph depicting the RTD ratio *(met22Δ/MET22^+^)* for *SUP4_oc_* variants, derived from the average % tY measured in *met22Δ* and *MET22*^+^ strains for each variant. Error bars represent propagated error for the quotient of *met22Δ/MET22^+^* average % tY values.

To directly determine if the similar *in vivo* RTD sensitivity of anticodon stem variants and acceptor stem variants was due to similar 5’ end accessibility, we measured 5’-3’ exonuclease activity *in vitro* on *SUP4_oc_* RTD variants, as we had previously done with tS(CGA) variants and Xrn1 (48). To assay decay, we used a biotinylated oligomer to purify *SUP4_oc_* variants together with endogenous tY(GUA), incubated the tRNAs with purified preparations of Xrn1 or Rat1/Rai1 complex (63,64), and assayed decay by poison primer extension, using the endogenous purified tY(GUA) as a control.

With Xrn1, we found that the U_2_C acceptor stem variant was highly susceptible to decay at 37°C, whereas the A_29_U and A_31_U anticodon stem variants were nearly as resistant as the negative control *SUP4_oc_* (Figure 2A and B). This assay was done at 37°C, as many RTD substrates are more sensitive to decay *in vivo* at high temperature (20,55). At 30°C, which is expected to be more stringent for tRNA decay, we still observed substantial sensitivity of the U_2_C variant to Xrn1, whereas the anticodon stem variants were both highly resistant (Supplementary Figure 1).

**Figure 2.**
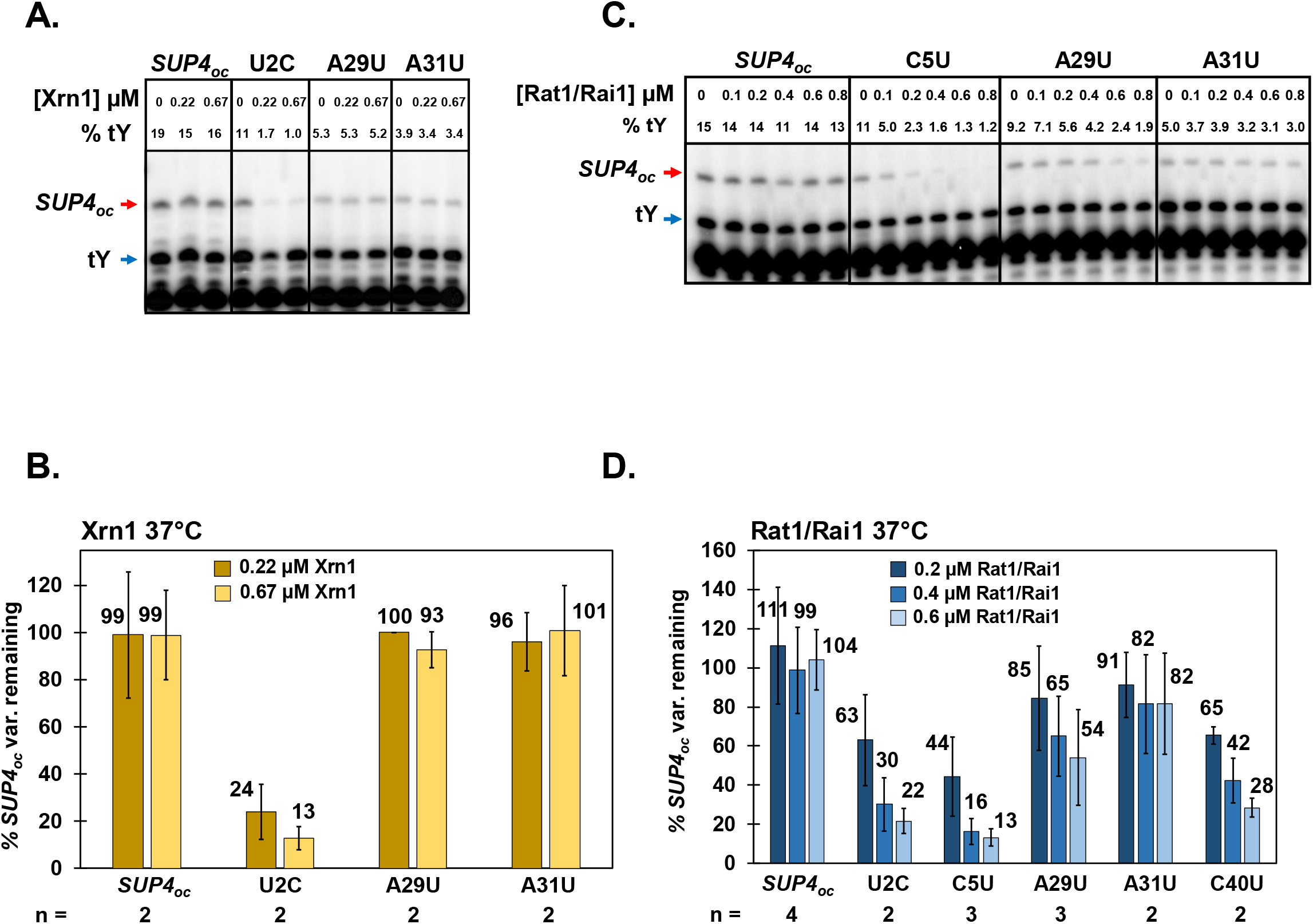
Anticodon stem *SUP4_oc_* variants are more resistant to 5’-3’ degradation than acceptor stem variants *in vitro*, despite their similar *in vivo* susceptibility. **(A) Analysis of degradation of *SUP4_oc_* variants by purified Xrn1 at 37°C.** Purified mixtures of *SUP4_oc_* variants and tY(GUA) were incubated with purified Xrn1 at indicated concentrations at 37°C for 20 minutes, Decay was analyzed by measuring the remaining *SUP4_oc_* variant tRNA, relative to tY(GUA), using poison primer extension analysis with primer P5 (nt 57-37) in the presence of ddCTP, producing a stop at G_34_ for endogenous tY(GUA) (blue arrow) and at G_30_ for *SUP4_oc_* variants (red arrow). **(B) Anticodon stem variants are more resistant than an acceptor stem variant to *in vitro* degradation by Xrn1 at 37°C.** The chart shows quantification of degradation of *SUP4_oc_* variants by purified Xrn1 at 37°C, analyzed at two different concentrations on each of two different days (n = 2). **(C) Analysis of degradation of *SUP4_oc_* variants by purified Rat1/Rai1 at 37°C.** Purified mixtures of *SUP4_oc_* variants and tY(GUA) were incubated with purified Rat1/Rai1 at indicated concentrations at 37°C for 1 hour, and decay was analyzed as in (A). **(D) Anticodon stem variants are more resistant than an acceptor stem variant to *in vitro* degradation by Rat1/Rai1 at 37°C.** The chart shows quantification of degradation by purified Rat1/Rai1 at 37°C, analyzed at three different concentrations on two-four different days (n = 2 − 4) as indicated.

With Rat1/Rai1 at 37°C, we observed very similar results to those with Xrn1. The *SUP4_oc_* acceptor stem U_2_C and C_5_U variants were highly susceptible to decay over a range of Rat1/Rai1 concentrations, whereas the A_31_U anticodon stem variant was highly resistant, and the A_29_U and C_40_U anticodon stem variants were moderately resistant (Figure 2C and D).

The finding that anticodon stem variants were consistently more resistant to 5’-3’ exonucleases *in vitro* than acceptor stem variants is consistent with our expectation that destabilization of the anticodon stem would have a limited effect on 5’ end exposure, but is seemingly contradictory with our *in vivo* data showing that anticodon stem and acceptor stem variants were similarly sensitive to RTD. The discrepancy between our *in vitro* and *in vivo* data implied that the degradation of anticodon stem variants by the RTD pathway might have an additional aspect *in vivo* that we were not able to account for *in vitro*.

### Anticodon stem variants accumulate pre-tRNA that is subject to RTD

We considered that the decay of anticodon stem variants measured *in vivo* might actually be the sum of the RTD pathway acting on both mature tRNA and pre-tRNA, although the RTD pathway had not previously been known to act on pre-tRNA. Since it was previously shown that an anticodon stem variant of another tRNA species had impaired tRNA maturation (65), it seemed reasonable that *SUP4_oc_* anticodon stem variants might accumulate pre-tRNA that could be targeted by the RTD pathway.

To determine if the pre-tRNAs of anticodon stem variants were also RTD substrates, we used poison primer extension to look for increased pre-tRNA in a *met22Δ* strain relative to a *MET22*^+^ strain, which would be diagnostic of pre-tRNA RTD. The primer for this experiment extended from nt 51 in the 3’ tRNA exon through the intron until residue intron-1 (In-1), and in principle would detect any of the unspliced precursors in the tRNA maturation pathway (Figure 3A).

**Figure 3.**
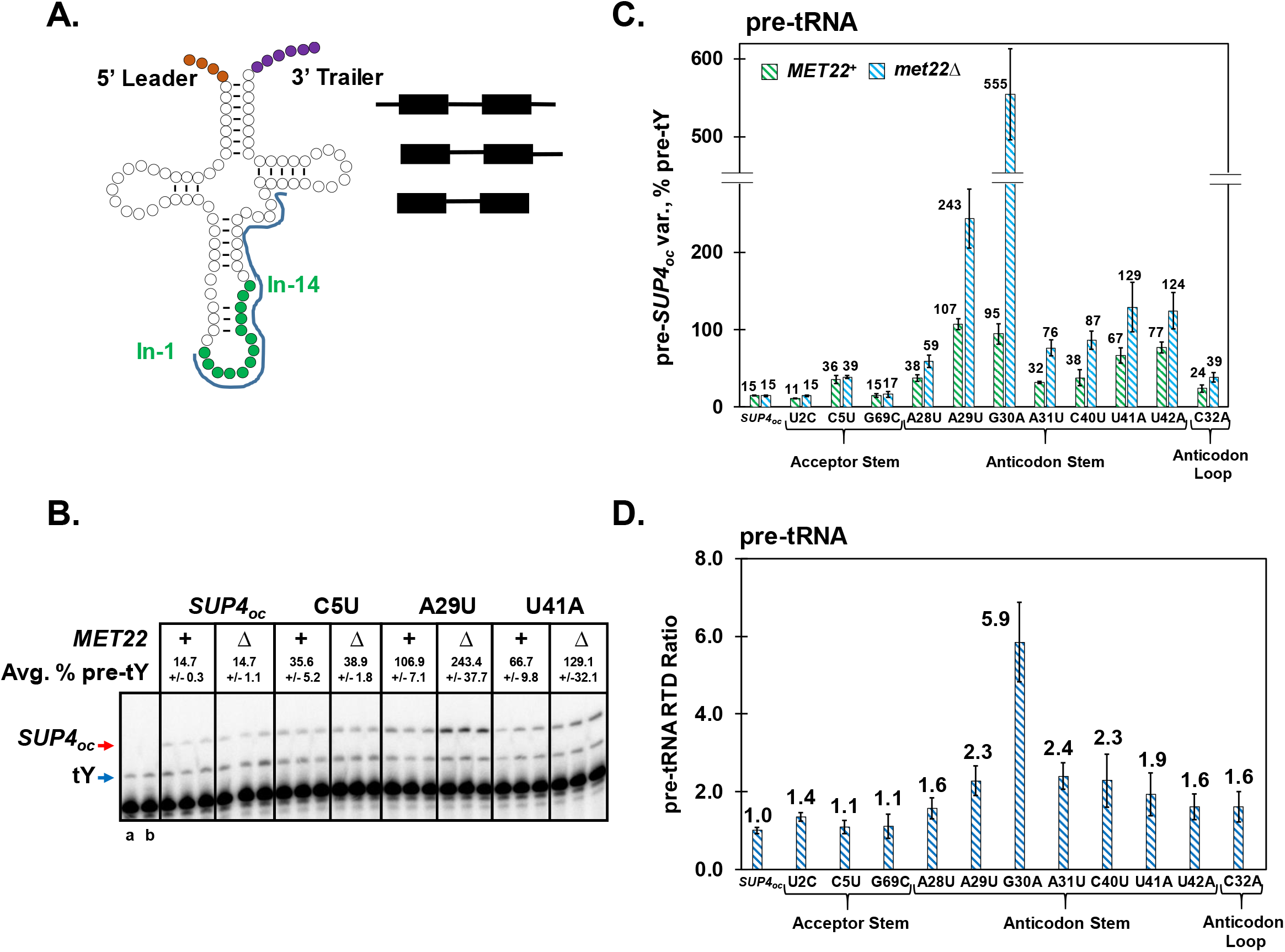
*SUP4_oc_* anticodon stem RTD variants are subject to pre-tRNA RTD, linked to pre-tRNA accumulation. **(A) Schematic of the precursors for tRNA^Tyr^.** The initial tRNA^Tyr^ transcript is depicted with a 5’ leader (orange), 3’ trailer (purple), and intron (green); primer MP 497 (51-[In-1]) used for measurement of pre-tRNA, blue line. Cartoons for pre-tY(GUA) intermediates are at the right; 5’, 3’ extended, initial transcript (top); 5’ mature 3’ extended (middle); and end matured (bottom) species. **(B) Poison primer extension analysis of levels of unspliced pre-tRNA of *SUP4_oc_* anticodon stem RTD variants.** Bulk RNA from strains with integrated *SUP4_oc_* variants was analyzed for pre-tRNA levels by poison primer extension as in Figure 1B, using primer MP 497 (51-[In-1]) in the presence of ddCTP, producing a stop at G_34_ for the endogenous pre-tY(GUA) (blue arrow) and a stop at G_30_ for *pre-SUP4_oc_* variants (red arrow). Controls: lane a, strain with no *SUP4_oc_* variant; lane b, strain with intronless *SUP4_oc_* variant. **(C) *SUP4_oc_* anticodon stem RTD variants have increased pre-tRNA in a *MET22*^+^ strain that is further increased in a *met22Δ* strain, compared to acceptor stem variants.** Bar chart depicting average % pre-tY(GUA) values with associated standard deviations for each of the analyzed variants (For *SUP4_oc_*, n=12 across four different experiments, for all other variants n=3); *MET22^+^*, green stripes, *met22Δ*, blue stripes. **(D) *SUP4_oc_* anticodon stem variants are subject to substantial pre-tRNA RTD, whereas acceptor stem variants are not.** Bar chart depicts the pre-tRNA RTD ratio *(met22Δ / MET22^+^)* for *SUP4_oc_* variants, derived from the % pre-tY(GUA) values in (C). Error bars represent propagated error for the quotient of *met22Δ/MET22^+^* average % pre-tY values.

We found evidence for substantial pre-tRNA RTD in our set of anticodon stem RTD variants. Initial experiments showed that both the *SUP4_oc_* A_29_U and U_41_A anticodon stem variants accumulated a large amount of pre-tRNA in a *MET22*^+^ strain relative to the accumulation of *SUP4_oc_* (7.1-fold and 4.5-fold respectively), and the pre-tRNA levels were further increased in a *met22Δ* strain relative to *SUP4_oc_* (16.2-fold and 8.6-fold), resulting in pre-tRNA RTD ratios of 2.3 and 1.9 respectively (Figure 3B,C, and D). By contrast, although the *SUP4_oc_* C_5_U acceptor stem variant accumulated a modest amount of pre-tRNA relative to *SUP4_oc_* (2.4-fold), there was no further increase in a *met22Δ* strain, resulting in a pre-tRNA RTD ratio of 1.1. These results extended to our entire set of anticodon stem RTD variants. Each of the five other anticodon stem RTD variants examined also showed significant accumulation of pre-tRNA relative to *SUP4_oc_* in *MET22*^+^ strains (2.1 fold or more), which in each case was further increased in a *met22Δ* strain, resulting in pre-tRNA RTD ratios ranging from 1.6 to 5.9 (Figure 3C and D). By contrast, the acceptor stem variants U_2_C and G_69_C showed no substantial accumulation of pre-tRNA and no significant pre-tRNA RTD ratio. Thus, it is a general property that *SUP4_oc_* anticodon stem RTD substrates have substantial increases in pre-tRNA levels in both *MET22*^+^ and *met22Δ* strains, and have substantial pre-tRNA RTD ratios.

This finding offers a simple explanation for the discrepancy between the similar RTD ratios of anticodon stem and acceptor stem variants *in vivo*, and the relative resistance of anticodon stem variants to 5’-3’ exonucleases *in vitro*. For anticodon stem variants, their susceptibility to pre-tRNA RTD would be captured by the poison primer extension measurements of mature *SUP4_oc_* tRNAs used to obtain RTD ratios (now referred to as overall RTD ratios), since overall RTD measured in this way is the sum of RTD of mature tRNA and of pre-tRNA. However, the reduced *in vitro* 5’-3’ exonuclease susceptibility of anticodon stem variants relative to acceptor stem variants would remain as a discriminating difference between the two classes of RTD variants.

It is not clear which nuclease is inhibited by the *met22Δ* mutation and is responsible for pre-tRNA RTD. Pre-tRNA RTD of *SUP4_oc_* A_29_U and U_41_A variants was not significantly inhibited by either a *rat1-107* mutation *or* an *xrn1Δ* mutation (Figure 4A, B, and C), each of which was previously found to significantly inhibit overall RTD in specific tRNA species (20,48). Similarly, we did not observe inhibition of pre-tRNA RTD with *trf4Δ* or *rrp6Δ* mutations (Figure 4D), each known to inhibit 3’-5’ pre-tRNA decay by the nuclear surveillance pathway (18,47), but not anticipated to be sensitive to the pAp buildup in a *met22Δ* mutant.

**Figure 4.**
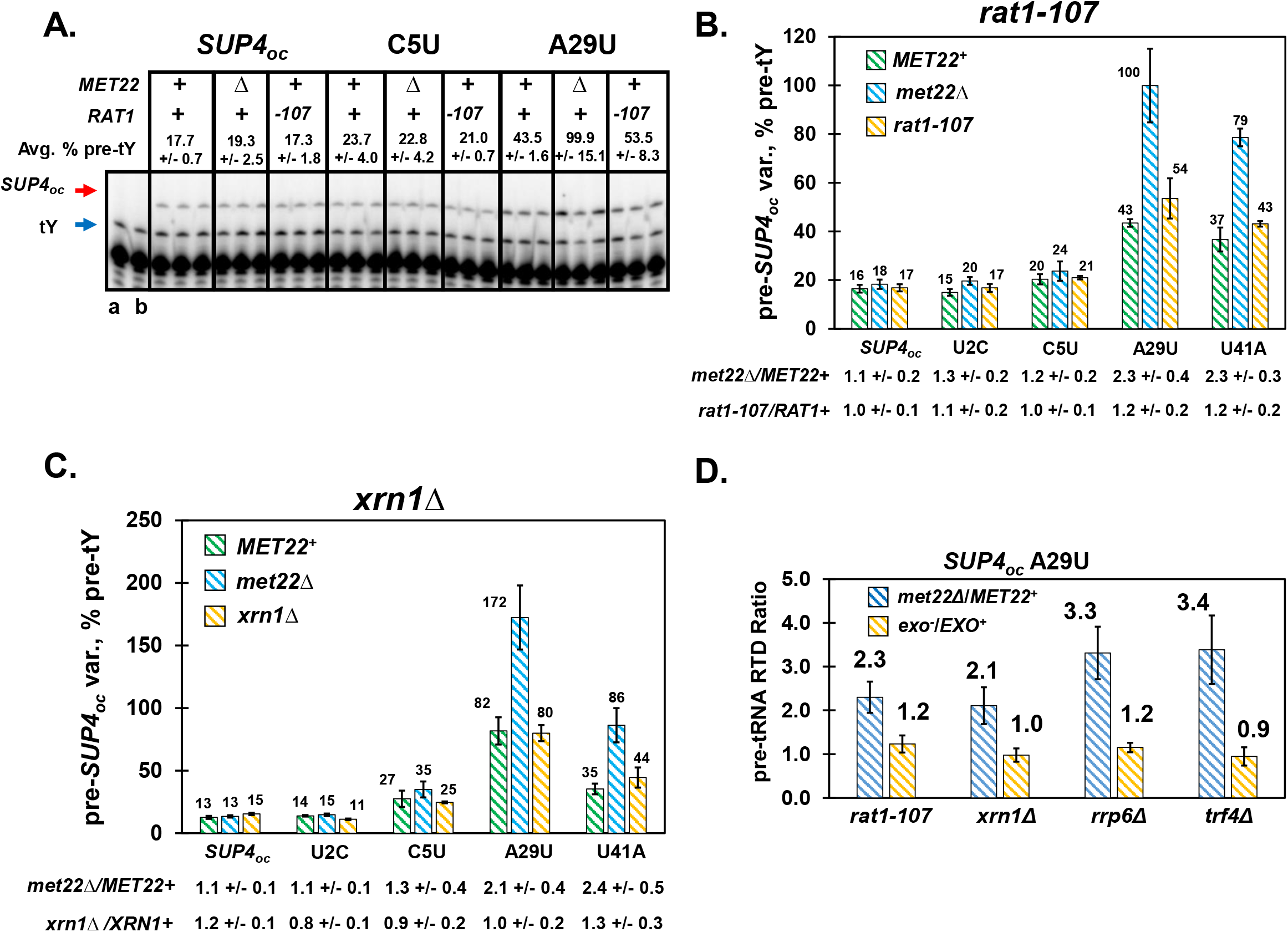
pre-tRNA RTD is not significantly inhibited by exonucleases known to mediate tRNA decay. **(A) Poison primer extension analysis of unspliced pre-tRNA in *MET22^+^, met22Δ*, and *rat1-107* strains.** RNA from *MET22^+^, met22Δ*, and *rat1-107* strains with *SUP4_oc_* variants as indicated was analyzed by poison primer extension for pre-tRNA levels as described in Figure 3B using the MP 497 primer (51-[In-1]) in the presence of ddCTP; endogenous pre-tY(GUA) (G_34_, blue arrow); *pre-SUP4_oc_* variants (G_30_, red arrow). lane a, no *SUP4_oc_* variant; lane b, intronless *SUP4_oc_* variant. **(B) The *rat1-107* mutation does not significantly suppress pre-tRNA RTD.** Bar chart depicting *pre-SUP4_oc_* variant levels (as % pre-tY) for variants in *MET22*^+^ (green stripes), *met22Δ* (blue stripes), and *rat1-107* (gold stripes) strains. Fold suppression is indicated below for the RTD ratio *(met22Δ/ MET22^+^)* or the corresponding *rat1-107/RAT1+* ratio. **(C) The *xrn1Δ* mutation does not significantly suppress pre-tRNA RTD.** Bar chart depicting *pre-SUP4_oc_* variant levels (as % pre-tY) for variants in *MET22*^+^ (green stripes), *met22Δ* (blue stripes), and *xrn1Δ* (gold stripes) strains. Fold suppression is indicated below for the RTD ratio *(met22Δ/ MET22^+^)* or the corresponding *xrn1 A /XRN1^+^* ratio. (D) pre-tRNA RTD of *SUP4_oc_* A_29_U is not suppressed by mutation of known exonucleases of the RTD pathway or known components of the nuclear surveillance pathway. For each mutant in the RTD pathway or the nuclear surveillance pathway, the pre-tRNA RTD ratio (blue stripes) is directly compared to the corresponding mutant ratio (gold stripes) to evaluate the effect of each mutation on pre-tRNA decay. Error bars represent propagated error for the indicated quotients of average % pre-tY values.

### Destabilization of anticodon-intron base pairing causes pre-tRNA accumulation and decay

The large accumulation of unspliced pre-tRNA associated with pre-tRNA RTD of *SUP4_oc_* anticodon stem variants suggested that perturbation of the pre-tRNA structure might in some way inhibit maturation to trigger the decay. In intron-containing pre-tRNAs, the anticodon forms part of a central helix with residues in the intron, surrounded by two bulges that comprise a bulge-helix-bulge region, which is recognized and cleaved by the splicing endonuclease in the first step of splicing (66–70). This characteristic pre-tRNA structure of the anticodon stem-loop and intron is predicted to be somewhat unstable in pre-tY(GUA) (62), which has only four base pairs in the central helix between the anticodon region and the intron (Figure 5A); moreover, the G_34_U *ochre* mutation of *SUP4_oc_* is predicted to destabilize the anticodon-intron pairing by disrupting the normal G_34_ pairing with intron residue In-11 (Fig. 5A), and to alter the pre-tRNA structure in the region. Thus, it seemed plausible that the U_34_ *ochre* mutation could be contributing to the accumulation of *SUP4_oc_* pre-tRNAs observed in some of the anticodon stem RTD variants.

**Figure 5.**
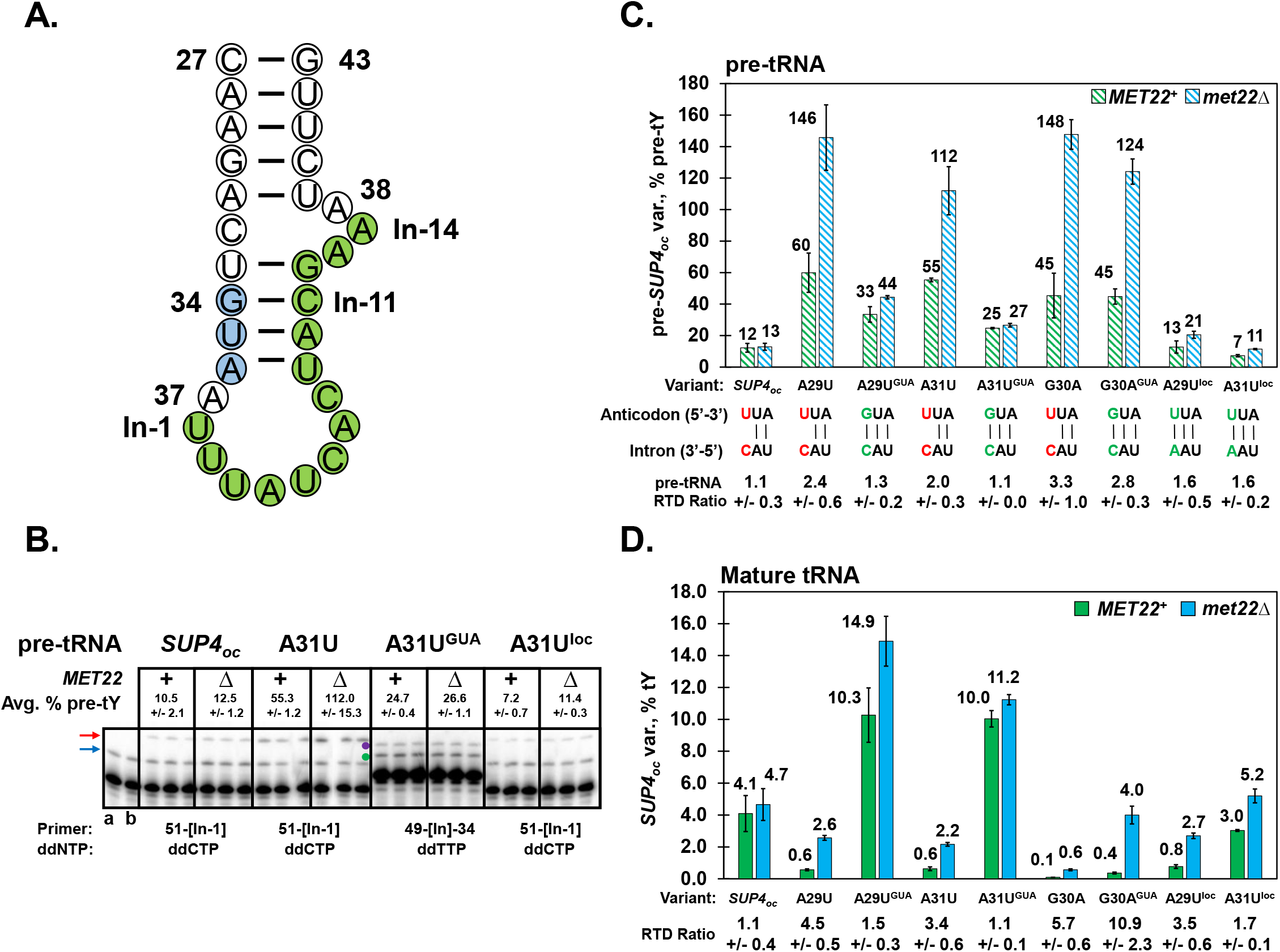
Anticodon-intron base pairing affects pre-tRNA accumulation and pre-tRNA RTD. **(A) Schematic of the anticodon stem-intron structure of pre-tRNA^Tyr^.** Nucleotides N27-N43 of the anticodon stem-loop are shown, as well as the intron (residues In-1 to In-14) arranged in the bulge-helix-bulge secondary structure common to substrates of the SEN complex. Anticodon, blue; intron, green. Note that *SUP4_oc_* has a G_34_U mutation. **(B) Analysis of pre-tRNA levels of a *SUP4_oc_* A_31_U variant with mutations predicted to alter pre-tRNA structure.** Poison primer extension was done as described in Figure 3B, using primer MP 497 (51-[In-1]) in the presence of ddCTP, producing a stop at G_34_ for endogenous pre-tY(GUA) (blue arrow) and at G_30_ for pre-tRNA of *SUP4_oc_*, the *SUP4_oc_* A_31_U variant and the *SUP4_oc_* A_31_U^I-oc^ variant (red arrow). The A_31_U^GUA^ variant was analyzed with primer MP 538 (49-[In]-34) in the presence of ddTTP, producing a stop at A31 for endogenous pre-tY(GUA) (green dot) and at A_29_ for the pre-A_31_U^GUA^ variant (purple dot). **(C) Restoration of anticodon-intron pairing suppresses pre-tRNA RTD for the *SUP4_oc_* A_29_U and A_31_U variants, but not the G_30_A variant.** Bar graph of % pre-tY levels for variants that have either mismatches (red) or pairing (green) between N_34_ and In-11. pre-tRNA RTD ratios (*met22Δ/MET22^+^*) for each variant are indicated in the bottom row. It should be noted that the measured % pre-tY values for the I-oc variants are artificially reduced due to a hybridization mismatch between the primer MP 497 (51-[In-1]) and the In-11 C-A mutation. **(D) Analysis of mature tRNA levels of *SUP4_oc_* variants indicates that pre-tRNA RTD can provide a substantial portion of the overall RTD ratio.** Overall RTD ratios for each variant are indicated at the bottom.

We tested the importance of the G_34_U *ochre* mutation in pre-tRNA RTD of three *SUP4_oc_* variants (A_29_U, G_30_A, and A_31_U) by reconstructing derivatives with a native GUA anticodon (A_29_U^GUA^ [ie, tY(GUA) A_29_U], G_30_A^GUA^, and A_31_U^GUA^) and comparing their pre-tRNA levels and pre-tRNA RTD ratios by poison primer extension.

The pre-tRNA RTD ratios were dramatically reduced in the A_29_U^GUA^ and A_31_U^GUA^ anticodon stem variants compared to the corresponding *SUP4_oc_* variants, from 2.4 to 1.3 for the A_29_U variant, and from 2.0 to 1.1 for the A_31_U variant (Figure 5B and C). Moreover, pre-tRNA accumulation was also dramatically reduced in these variants in the *met22Δ* strain, 3.3-fold in the A_29_U variant (from 146 to 44 % pre-tY) and 4.1 fold in the A_31_U variant. In contrast, the pre-tRNA RTD ratio was only slightly reduced for the G_30_A^GUA^ variant compared to the *SUP4_oc_* G_30_A variant, from 3.3 to 2.8, and the pre-tRNA accumulation was only slightly reduced in the *met22Δ* strain (1.2-fold). The results with all three tY(GUA) variants are consistent with a model in which pre-tRNA RTD is driven by pre-tRNA accumulation, which is in turn due to perturbation of the pre-tRNA structure that is important for splicing. The persistence of the G_30_A mutation in causing pre-tRNA RTD with either a native GUA anticodon or an *ochre* anticodon is consistent with the greater predicted disruption in pre-tRNA structure caused by the G_30_A mutation, compared to either the A_29_U or A_31_U mutations (62).

This analysis also revealed two other important properties of pre-tRNA RTD. First, pre-tRNA RTD comprised a major component of the overall RTD of two of the variants, since the dramatic reduction in pre-tRNA RTD ratios for the A_29_U^GUA^ and A_31_U^GUA^ was matched by a similar reduction in their overall RTD ratios (Figure 5D). Second, substantial amounts of pre-tRNA RTD can occur in anticodon stem RTD variants with a natural GUA anticodon, since the G_30_A^GUA^ variant had a large pre-tRNA RTD ratio of 2.8.

We further tested the importance of anticodon-intron pairing in pre-tRNA RTD by making a compensatory mutation of In-11 to restore the pairing of *SUP4_oc_* residue U34 (making a *SUP4_oc_* variant denoted as ^I-oc^), albeit with a U-A pair instead of the native G-C pair (Fig. 5A). We found that the resulting *SUP4_oc_* A_29_U^I-oc^ and *SUP4_oc_* A_31_U^I-oc^ variants each had the anticipated reduction in pre-tRNA RTD ratio relative to the corresponding *SUP4_oc_* variants, but not as much of a reduction as in the corresponding tY^GUA^ variants. Consistent with an intermediate effect on pre-tRNA RTD, we observed an intermediate reduction in overall RTD for the A_29_U^I-oc^ and A_31_U^I-oc^ variants relative to their *SUP4_oc_* and tY^GUA^ counterparts (Figure 5D). The observed correspondence between restoration of anticodon-intron pairing and reduced pre-tRNA RTD emphasizes the importance of the structure of the pre-tRNA in the region of the anticodon stem-loop and intron for pre-tRNA RTD.

### Anticodon stem mutations cause both RTD and pre-tRNA RTD in *tS(CGA)*

The finding of RTD associated with anticodon stem variants led us to speculate that anticodon stem RTD variants would be found in other intron-containing tRNAs, and that pre-tRNA RTD would likely be associated with them. We therefore attempted to design additional anticodon stem variants that would be subject to RTD in *tS(CGA)* (Figure 6A), a single copy essential tRNA gene that could be examined for function by a simple genetic test (48). We integrated a *tS(CGA)* anticodon stem variant into a *tS(CGA)Δ[URA3 tS(CGA)]* strain, so that cell growth dependent on the integrated variant could be tested after FOA selection to remove the *URA3*^+^ plasmid bearing the wild type *tS(CGA)* gene. We found that strains bearing each of three tested anticodon stem *tS(CGA)* variants (A_29_U, C_40_U, and C_42_U) behaved like RTD substrates, because they exhibited temperature sensitivity that could be suppressed by a *met22Δ* mutation (Figure 6A and B), as was observed for the control *tan1Δ trm44Δ* mutant, previously shown to be due to RTD (20,71).

**Figure 6.**
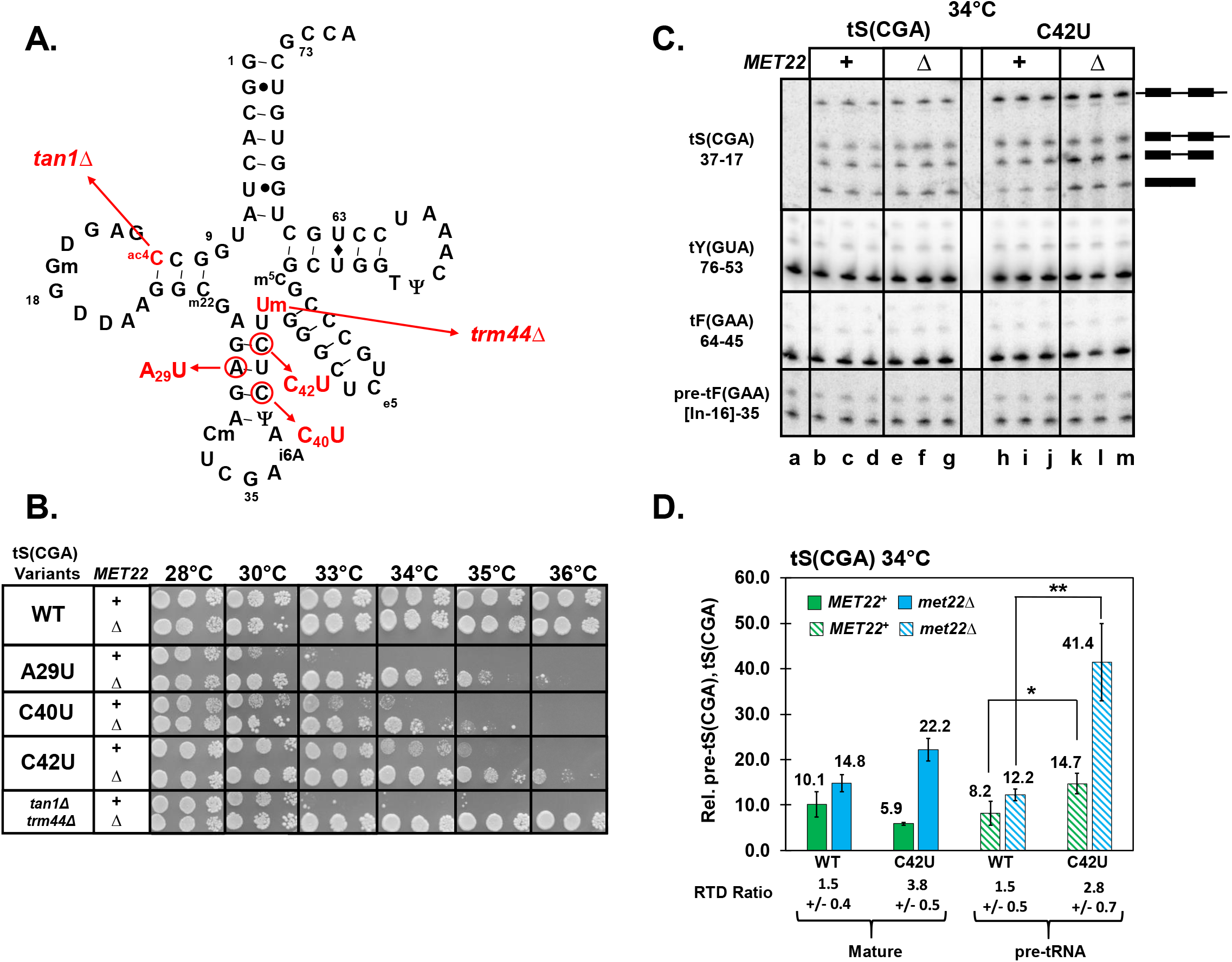
Anticodon stem variants of *tS(CGA)* provoke RTD, accompanied by pre-tRNA RTD for the *tS(CGA)* C_42_U variant. **(A) Schematic of mature *tS(CGA)* and variants analyzed.** Variants and modification mutants tested are indicated in red. **(B) Genetic analysis demonstrates that several *tS(CGA)* anticodon stem variants provoke RTD.** *MET22*^+^ and *met22Δ* strains with a *tS(CGA)A* mutation and a single integrated *tS(CGA)* variant as indicated were tested for growth by plating serial dilutions on YPD media, and incubating plates at the temperatures indicated. **(C) Northern analysis of *tS(CGA)* and *tS(CGA)* C_42_U tRNA levels in *MET22*^+^ and *met22Δ* strains grown at 34°C.** Strains with a *tS(CGA)Δ* mutation, an integrated tS(CGA) variant, and a [*tS(GCU/CGA) LEU2 CEN]* plasmid (to allow for growth at a non-permissive temperature), were grown at 34°C, and bulk RNA was analyzed by Northern analysis using probes with target regions as indicated. Lane a, strain with no integrated *tS(CGA)* variant, for hybridization control. Pre-tRNA and mature tRNA species indicated by cartoons. **(D) The *tS(CGA)* C_42_U variant is subject to RTD and pre-tRNA RTD.** Pre-tS(CGA) and mature tS(CGA) were quantified relative to hybridization of tY (x 100) to control for loading, and RTD and pre-tRNA RTD ratios are indicated at the bottom. The statistical significance of pre-tRNA accumulation was evaluated using a 1-tailed Student’s t-test assuming equal variance.

To assess tRNA decay of these variants, we first introduced a plasmid containing a tS(GCU/CGA) hybrid tRNA, bearing the body of tS(GCU) and a CGA anticodon, into the *tS(CGA)Δ* strain with an integrated *tS(CGA)* variant, thus allowing growth of strains at non-permissive temperatures and facile measurement of tS(CGA) variant levels by northern analysis. To probe specifically for *tS(CGA)* and not the nearly identical isoacceptor *tS(UGA)*, we used a probe that hybridized to nt 17-37 of tS(CGA), which differs from tS(UGA) at residues 28 and 34, and used hybridization conditions yielding no signal in a WT strain that contained only tS(GCU/CGA) as a source of tS(CGA) (Figure 6C, lane a). At 34°C, the *tS(CGA)* C_42_U variant had a substantial overall RTD ratio of 3.8, accompanied by a significant accumulation of unspliced end-matured pre-tRNA and a pre-tRNA RTD ratio of 2.8 (Figure 6C and D). There was no corresponding *met22Δ-dependent* accumulation of either the primary transcript or the 5’ end processed unspliced intermediate. At 32°C, the *tS(CGA)* C_40_U variant also had a substantial overall RTD ratio of 3.9 and a distinct pre-tRNA RTD ratio of 1.8, albeit accompanied by a modest p-value of 0.065 for pre-tRNA accumulation in the *met22Δ* background (Supplementary Figure 2A and B). By contrast, although the *tS(CGA)* A_29_U variant had substantial overall RTD, there was no obvious accumulation or degradation of pre-tRNA (Supplementary Figure 2C and D). These results provide compelling evidence for overall RTD of anticodon stem variants of *tS(CGA)*, accompanied in two cases by pre-tRNA RTD and accumulation of unspliced pre-tRNA. Thus, the pre-tRNA RTD decay pathway is a general phenomenon, and for the *tS(CGA)* C_42_U variant and the *tS(CGA)* C_40_U variant, pre-tRNA RTD occurred after end processing and before splicing.

### Inhibition of splicing exacerbates pre-tRNA RTD of the *SUP4_oc_ A_29_U* variant, whereas inhibition of nuclear export provokes nuclear surveillance

To precisely define the specific tRNA maturation defect of *SUP4_oc_* anticodon stem variants that leads to pre-tRNA RTD and the associated accumulation of unspliced pre-tRNA, we used mutants that perturbed specific steps of maturation. In yeast, intron-containing pre-tRNAs are end-processed in the nucleus (21), exported to the cytoplasm by either *LOS1* or the *MEX67/MTR2* heterodimer (72), and then spliced on the surface of the mitochondria by the SEN complex (24,25). Since our analysis of the *tS(CGA)* C_42_U variant suggested that pre-tRNA RTD occurred after end processing, we focused on the nuclear export and splicing steps for analysis of pre-tRNA RTD of *SUP4_oc_* variants. If the pre-tRNA of *SUP4_oc_* anticodon stem variants was degraded at one of these steps, then slowing down that step would increase the availability of the pre-tRNA at that stage, resulting in increased pre-tRNA RTD.

To examine the importance of the nuclear export step in pre-tRNA RTD, we used the well-studied *los1Δ* mutation, which causes nuclear accumulation of intron-containing pre-tRNA (73,74); and to examine the splicing step, we used a temperature sensitive *sen2-42* allele of the Sen2 catalytic subunit of the SEN complex, which accumulates unspliced pre-tRNA at 32°C (25). We generated *los1Δ* and *sen2-42* mutants in both a *MET22*^+^ and *met22Δ* background, and then integrated the *SUP4_oc_* A_29_U anticodon stem variant to analyze pre-tRNA RTD by Northern analysis. As we had observed for the tS(CGA) C_42_U variant, analysis of the control *LOS1*^+^ and *SEN2*^+^ strains showed that pre-tRNA RTD of the *SUP4_oc_* A_29_U variant occurred at the level of end-matured unspliced pre-tRNA, rather than at the level of primary transcript or 5’-trimmed 3’-trailer containing unspliced pre-tRNA (Fig. 7A, lanes b-g and o-t),.

**Figure 7.**
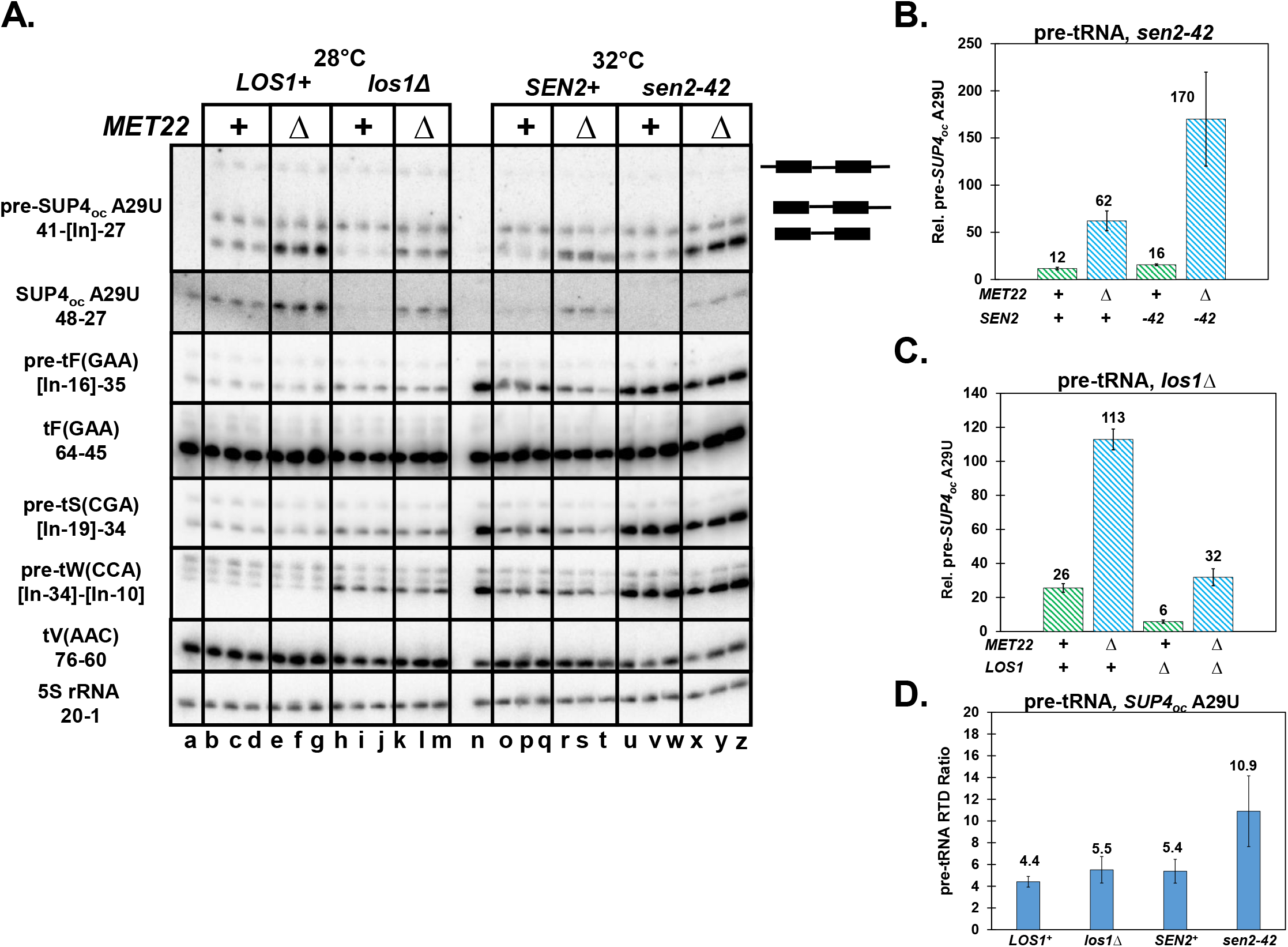
Pre-tRNA RTD of the *SUP4_oc_* A_29_U variant is exacerbated by inhibition of splicing, but not by inhibition of nuclear export. **(A) Northern analysis of *SUP4_oc_* A_29_U variant tRNA and pre-tRNA levels in *los1Δ* and *sen2-42* mutant strains in a *MET22*^+^ or *met22Δ* background.** Left side: *SUP4_oc_* A_29_U LOS1^+^ and *SUP4_oc_ A_29_U los1Δ* strains with *MET22*^+^ or *met22Δ* mutations as indicated were grown at 28°C, and tRNA species were analyzed by Northern analysis with probes as indicated. Lane a, a *MET22*^+^ strain with no integrated *SUP4_oc_* variant. Right side: *SUP4_oc_ A_29_U sen2Δ [SEN2^+^ TRP1 CEN]* and *SUP4_oc_ A_29_U sen2A [sen2-42 TRP1 CEN]* strains were grown at 32°C and analyzed for tRNA species as for left side; Lane n, a *sen2-42* strain with no integrated *SUP4_oc_* variant. **(B) Quantification of pre-tRNA levels for the *SUP4_oc_* A_29_U variant in *sen2-42* or *SEN2+* strains with a *MET22*^+^ or *met22Δ* mutation.** Pre-tRNA levels were quantified relative to hybridization of 5S to control for loading. **(C) Quantification of pre-tRNA levels for the *SUP4_oc_ A_29_U* variant in *LOS1*^+^ and *los1Δ* strains with a *MET22*^+^ or *met22Δ* mutation.** Pre-tRNA levels were quantified as in (B). **(D) Comparison of pre-tRNA RTD ratios for the *SUP4_oc_ A_29_U* variant in *los1Δ* and *sen2-42* strains relative to control WT strains.** Error bars represent propagated error for the indicated quotients of average % pre-tY values.

Our analysis of the *sen2-42* strain showed that inhibition of splicing exacerbated pre-tRNA RTD of the *SUP4_oc_* A_29_U anticodon stem variant. As expected because of the inhibition of splicing in the *sen2-42* mutant, there was a large increase in intron-containing pre-tRNA for each of three tested tRNAs (pre-tF(GAA), pre-tS(CGA), and pre-tW(CCA)), but none of these pre-tRNAs had significant pre-tRNA RTD (Figure 7A, Supplementary Figure 3A). By contrast, there was a significant increase in the pre-tRNA RTD ratio of the *SUP4_oc_* A_29_U anticodon stem variant in the *sen2-42* mutant relative to that in the *SEN2*^+^ strain, from 5.4 to 10.9 (Figure 7B and D). This result provides strong evidence that the maturation defect of the *SUP4_oc_* A_29_U variant that causes pre-tRNA RTD lies at or before the splicing step.

As expected if pre-tRNA RTD occurs after export from the nucleus, inhibition of pre-tRNA nuclear export by the *los1Δ* mutation did not affect pre-tRNA RTD of the *SUP4_oc_* A_29_U variant. As anticipated for impaired pre-tRNA export, pre-tRNA levels for each of three controls (pre-tF(GAA), pre-tS(CGA) and pre-tW(CCA)) were significantly increased in both a *los1Δ MET22*^+^ and a *los1Δ met22Δ* strain, relative to the corresponding *LOS1*^+^ control strains (Figure 7A, Supplementary Figure 3B). By contrast, we found that the levels of *SUP4_oc_* A_29_U pre-tRNA were unexpectedly decreased in the *los1Δ* strain (compared to the *LOS1*^+^ control strain), and were decreased to similar extents in *MET22*^+^ and *met22Δ* strains, resulting in a very similar pre-tRNA RTD ratio in both a *LOS1*^+^ and a *los1Δ* background (Fig. 7A, compare lanes h-m to b-g, Figure 7C and D). Thus, the reduced pre-tRNA levels of the *SUP4_oc_* A_29_U variant in a *los1Δ* mutant appear to be due to another decay pathway, not regulated by *MET22*.

This additional pre-tRNA decay observed in a *los1Δ* mutant is due to the nuclear surveillance pathway, since the reduced pre-tRNA levels of the *SUP4_oc_* A_29_U variant in a *los1Δ* strain, relative to a *LOS1*^+^ strain, were restored upon mutation of a known component of the nuclear exosome, *RRP6*, (Figure 8A and B), which degrades substrate pre-tRNAs from the 3’ end (75,76). The concomitant increase in mature *SUP4_oc_* A_29_U tRNA levels in the *los1Δ rrp6Δ* strain (Figure 7A) underscores that pre-tRNA decay by the nuclear surveillance pathway contributes significantly to overall production of mature tRNA for this variant in this background.

**Figure 8.**
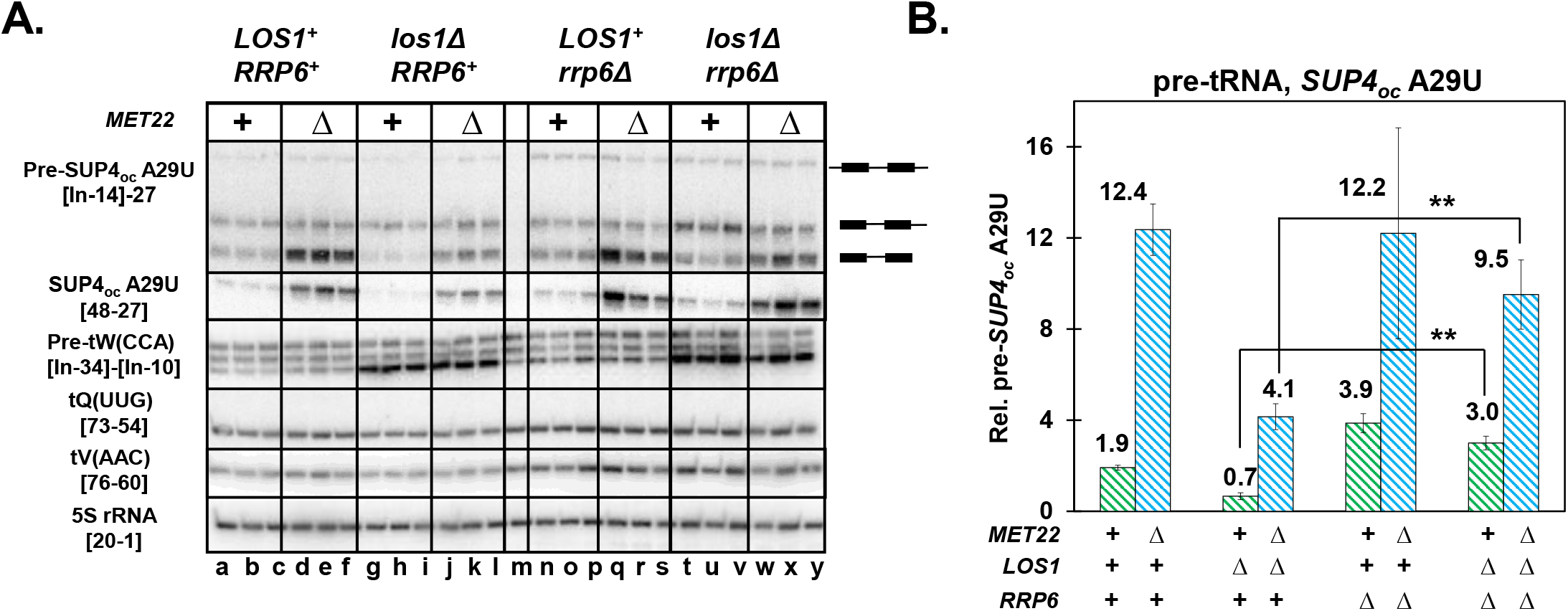
The nuclear surveillance pathway can degrade pre-tRNA of a *SUP4_oc_* A_29_U variant in a *los1Δ* strain. **(A) Northern analysis of the effect of an *rrp6Δ* mutation on pre-tRNA decay of a *SUP4_oc_* A_29_U variant in a *los1Δ* strain.** *SUP4_oc_* A_29_U *LOS1*^+^ and *SUP4_oc_* A_29_U *los1Δ* strains with *MET22*^+^ or *met22Δ* mutations and *RRP6*^+^ or *rrp6Δ* mutations as indicated were grown at 28°C, and tRNA species were analyzed by Northern analysis with probes as indicated. m, a *LOS1^+^ MET22^+^ rrp6Δ* strain with no integrated *SUP4_oc_* variant. **(B) Quantification of the effect of an *rrp6Δ* mutation on the pre-tRNA decay of the *SUP4_oc_* A_29_U variant in a *los1Δ* strain.** Pre-tRNA levels were quantified relative to 5S as in Figure 7B. The statistical significance of pre-tRNA increase between *los1Δ* and *los1Δ rrp6Δ* was evaluated using a 1-tailed Student’s t-test assuming equal variance.

## DISCUSSION

The results described here show that RTD of anticodon stem variants occurs in part through a previously undescribed pre-tRNA RTD pathway, which targets pre-tRNAs that accumulate due to altered structure of the anticodon stem-loop and intron, likely resulting in inefficient splicing. Pre-tRNA RTD can contribute substantially to overall RTD since mutations that suppress pre-tRNA RTD result in comparable suppression of overall RTD (Figure 5), and pre-tRNA RTD is a general phenomenon, since it can occur in both tY(GUA) and tS(CGA) variants bearing native anticodons and thus participating in normal translation (Figures 5 and 6). In all previously described cases, RTD has only been documented to act on mature tRNA (19,20,49), and decay of pre-tRNA has only been attributed to 3’-5’ exonucleolytic decay by the nuclear surveillance pathway (17,18,47).

The accumulation of pre-tRNA that drives pre-tRNA RTD appears to be directly related to the pre-tRNA anticodon stem-loop and intron structure, which is important for splicing *in vivo*. Anticodon-intron pairing is highly conserved in archaeal and yeast intron-containing pre-tRNAs as part of the bulge-helix-bulge motif important for splicing (66,67,69,70,77–79), and may also be important in metazoans (68). The importance of the central helix is underscored by our results that disruption of the central helix tended to increase pre-tRNA RTD, whereas restoration of the helix tended to reduce pre-tRNA RTD (Figure 5). The importance of the 3’ bulge structure is underscored by prominent pre-tRNA RTD in variants predicted to have an intact central helix, but with mutations affecting the bulge, including the *tY(GUA)* G_30_A, *tS(CGA)* C_40_U, and *tS(CGA)* C_42_U variants (Figures 5 and 6).

Remarkably, the pre-tRNA RTD pathway is not inhibited by mutation of the known exonucleases of the RTD pathway, or of known components of the nuclear surveillance pathway (Figure 4). We envision three possibilities for the mechanism. First, pre-tRNA RTD might actually be due to Rat1, if the *rat1-107* allele is not strong enough to inhibit pre-tRNA RTD. Since *RAT1* is essential, obtaining such a mutation might be difficult, and use of the *rat1^ts^* allele (80,81) would not be possible for experiments at 28°C; furthermore, the effect of the *rat1-107* mutation on Rat1 activity has not been measured at 28°C. Second, pre-tRNA RTD might be due in part to another 5’-3’ exonuclease, such as Dxo1 or Rrp17 (82,83), which are expected to be inhibited by the pAp accumulation caused by a met22Δ mutation (51,52,84). Third, it is conceivable that pre-tRNA RTD might involve decay by an endonuclease such as Rny1 (85) or the splicing endonuclease itself (86), due to the prolonged presence of unspliced pre-tRNA as a consequence of the altered structure of the pre-tRNA. The involvement of an endonuclease in pre-tRNA decay would be similar in this regard to nonsense-mediated decay of mRNA, which involves both an endonuclease and an exonuclease (87), and would require inhibition by pAp to explain the met22Δ-dependent effects. Full understanding of the mechanism of pre-tRNA decay will undoubtedly lead to new insights in RNA quality control mechanisms.

The results outlined here suggest a model of opportunistic decay in which tRNA is constantly under surveillance through each step of biogenesis and as mature tRNA, with decay occurring as a consequence of the availability and instability of each state of the tRNA, by either the RTD pathway or the nuclear surveillance pathway. The nuclear surveillance pathway targets initial pre-tRNA transcripts of pre-tRNA_i_^Met^ lacking m^1^A_58_, and a large portion of WT tRNAs soon after transcription (17,18,47). Results described here show that end-matured unspliced pre-tRNA is subjected to two rounds of quality control: It is targeted for pre-tRNA RTD after nuclear export and before or at the splicing step, based on analysis of the effect on pre-tRNA RTD of mutations that inhibit each step (Figure 7); and it can be targeted by the nuclear surveillance pathway, particularly if it is retained in the nucleus. In this model, the substantial overall RTD of anticodon stem *SUP4_oc_* variants is due to a combination of pre-tRNA RTD, triggered by their altered intron-exon structure and presumed inefficient splicing, coupled with the distinct, but relatively inefficient, 5’-3’ decay of anticodon stem variants after splicing, resulting in similar overall RTD to that of acceptor stem variants. Subsequent to maturation, tRNAs with destabilizing mutations or lacking specific body modifications are targeted by the RTD pathway (19,20,49,54), particularly at elevated temperatures, which leads to increased instability and attack (48,55). RTD is in competition with cellular proteins that bind charged tRNA, including EF-1A (49,88) and tRNA synthetases such as ValRS for RTD of tV(AAC) (88). RTD may also be in competition with the trafficking machinery, since mature cytoplasmic tRNA lacking m^2,2^G is degraded by RTD or is subject to retrograde import into the nucleus (50). This model of opportunistic decay is similar to the previously proposed scavenger mechanism attributed to RNA quality control pathways that also monitor tRNAs in *Escherichia coli* (89–91).

This model of opportunistic decay underscores the high degree of evolutionary tuning required for all tRNA species to avoid the multiple quality control decay pathways, in addition to their more well studied evolution to function efficiently and accurately in translation. Since availability and instability are twin drivers of the RTD pathway, and likely also of the nuclear surveillance pathway, each tRNA species must have evolved for optimized processing efficiency of each intermediate, in addition to stability, to prevent access by the nucleases of the decay pathways, as well as to ensure timely maturation of tRNA for increased translation efficiency. Since the components of RTD and nuclear surveillance are widely conserved in eukaryotes, it seems likely that the tRNA decay pathways will be conserved in metazoans (53), and that the selection of tRNAs for processing efficiency and stability will also be conserved. This raises the possibility that some of the numerous isodecoders in metazoans (92,93) will have different susceptibilities to decay in addition to differences in expression and function (93,94).

## ACKNOWLEDGEM ENTS

We thank Elizabeth Grayhack for valuable discussions during the course of this work and comments on the manuscript.

---

**Supplementary Table 1.**
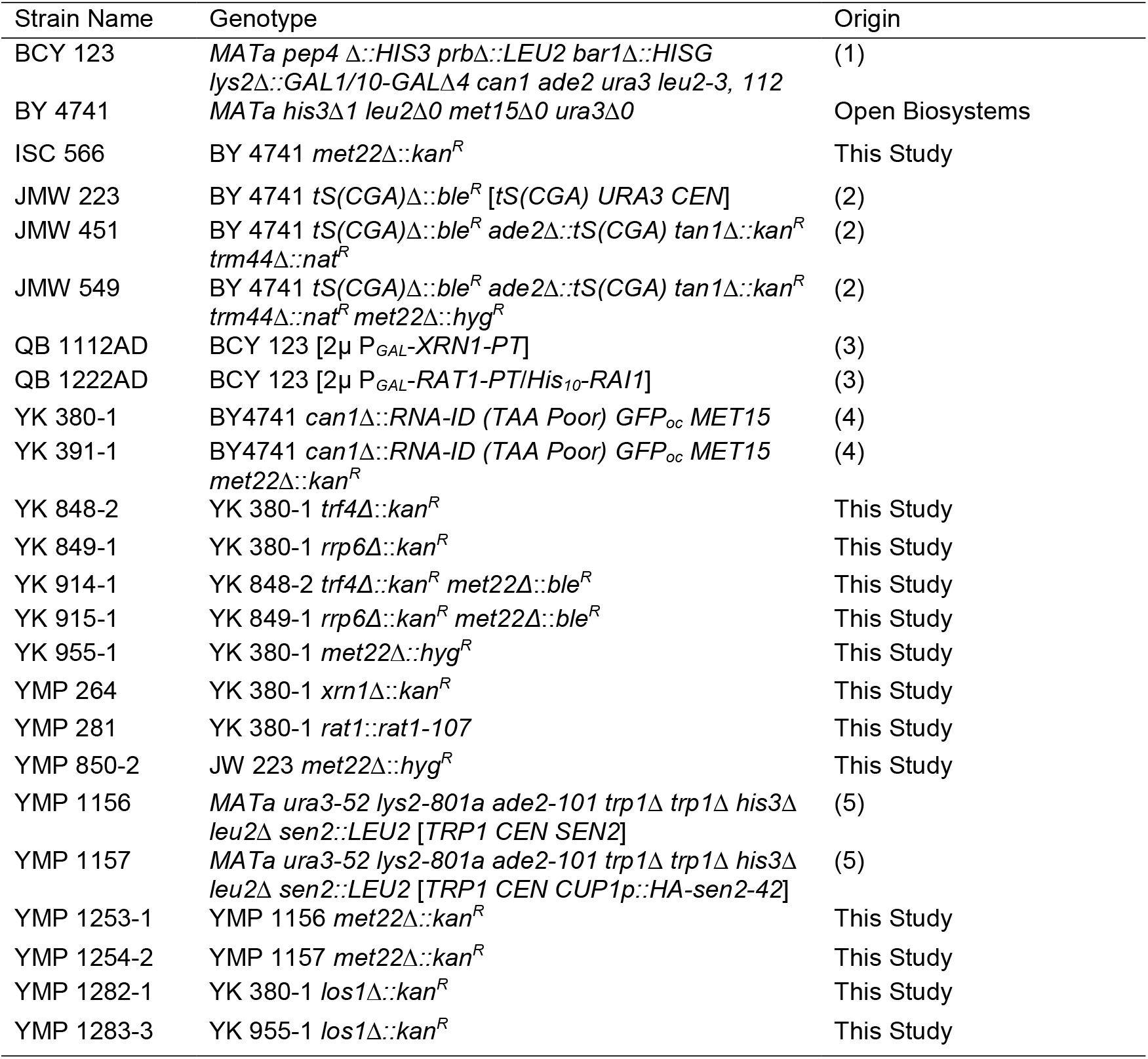
Strains used in this study

| Strain Name | Genotype | Origin |
| --- | --- | --- |
| BCY 123 | *MATa pep4 Δ::HIS3 prbΔ::LEU2 bar1Δ::HISG lys2Δ::GAL1/10-GALΔ4 can1 ade2 ura3 leu2-3, 112* | (1) |
| BY 4741 | *MATa his3Δ1 leu2Δ0 met15Δ0 ura3Δ0* | Open Biosystems |
| ISC 566 | BY 4741 *met22Δ::kan<sup>R</sup>* | This Study |
| JMW 223 | BY 4741 *tS(CGA)Δ::ble<sup>R</sup> [tS(CGA) URA3 CEN]* | (2) |
| JMW 451 | BY 4741 *tS(CGA)Δ::ble<sup>R</sup> ade2Δ::tS(CGA) tan1Δ::kan<sup>R</sup> trm44Δ::nat<sup>R</sup>* | (2) |
| JMW 549 | BY 4741 *tS(CGA)Δ::ble<sup>R</sup> ade2Δ::tS(CGA) tan1Δ::kan<sup>R</sup> trm44Δ::nat<sup>R</sup> met22Δ::hyg<sup>R</sup>* | (2) |
| QB 1112AD | BCY 123 [2μ P<sub>GAL</sub>-XRN1-PT] | (3) |
| QB 1222AD | BCY 123 [2μ P<sub>GAL</sub>-RAT1-PT/His<sub>10</sub>-RAI1] | (3) |
| YK 380-1 | BY4741 *can1Δ::RNA-ID (TAA Poor) GFP<sub>oc</sub> MET15* | (4) |
| YK 391-1 | BY4741 *can1Δ::RNA-ID (TAA Poor) GFP<sub>oc</sub> MET15 met22Δ::kan<sup>R</sup>* | (4) |
| YK 848-2 | YK 380-1 *trf4Δ::kan<sup>R</sup>* | This Study |
| YK 849-1 | YK 380-1 *rrp6Δ::kan<sup>R</sup>* | This Study |
| YK 914-1 | YK 848-2 *trf4Δ::kan<sup>R</sup> met22Δ::ble<sup>R</sup>* | This Study |
| YK 915-1 | YK 849-1 *rrp6Δ::kan<sup>R</sup> met22Δ::ble<sup>R</sup>* | This Study |
| YK 955-1 | YK 380-1 *met22Δ::hyg<sup>R</sup>* | This Study |
| YMP 264 | YK 380-1 *xrn1Δ::kan<sup>R</sup>* | This Study |
| YMP 281 | YK 380-1 *rat1::rat1-107* | This Study |
| YMP 850-2 | JW 223 *met22Δ::hyg<sup>R</sup>* | This Study |
| YMP 1156 | *MATa ura3-52 lys2-801a ade2-101 trp1Δ trp1Δ his3Δ leu2Δ sen2::LEU2 [TRP1 CEN SEN2]* | (5) |
| YMP 1157 | *MATa ura3-52 lys2-801a ade2-101 trp1Δ trp1Δ his3Δ leu2Δ sen2::LEU2 [TRP1 CEN CUP1p::HA-sen2-42]* | (5) |
| YMP 1253-1 | YMP 1156 *met22Δ::kan<sup>R</sup>* | This Study |
| YMP 1254-2 | YMP 1157 *met22Δ::kan<sup>R</sup>* | This Study |
| YMP 1282-1 | YK 380-1 *los1Δ::kan<sup>R</sup>* | This Study |
| YMP 1283-3 | YK 955-1 *los1Δ::kan<sup>R</sup>* | This Study |

**Supplementary Table 2.** Primers used in poison primer extension

| Name | Target RNA | Hybridization Region 5'-3' |
| --- | --- | --- |
| P5 | tY(GUA) | 57-37 |
| P7 | tY(GUA) | 62-43 |
| P9 | tY(GUA) | 57-33 |
| MP 497 | tY(GUA) | 51-[In-1] |
| MP 538 | tY(GUA) | 49-[In]-34 |
| MP 539 | tY(GUA) | 55-31 |
| MP 540 | tY(GUA) | 46-[In]-31 |

**Supplementary Table 3.** Oligomers used for northern analysis

| Target RNA | Name | Hybridization Region 5'-3' |
| --- | --- | --- |
| *SUP4<sub>oc</sub>* A29U | TDZ 231 | [48-27] |
| tF(GAA) | tRNAPheLoop | 64-45 |
| tQ(UUG) | 3'-tRNAGln | [73-54] |
| tS(CGA) | TDZ 199 | [e8-33] |
| tS(CGA) | MP 586 | 37-17 |
| tS(GCU) | Ser-(GCU)1 | e7-31 |
| tV(AAC) | Val_CCA | 76-60 |
| tW(CCA) | TrpP1 | 33-15 |
| tY(GUA) | tY(GUA) | 76-53 |
| pre-tF(GAA) | Phe GAA intron | [In-16]-35 |
| pre-tS(CGA) | MP 469 | [In-19]-34 |
| pre-tW(CCA) | TDZ 235 | [In-34]-[In-10] |
| pre-tY(GUA) | MP 564 | 43-[In-1] |
| pre-*SUP4<sub>oc</sub>* A29U | MP 567 | [In-14]-27 |
| pre-*SUP4<sub>oc</sub>* A29U | MP 524 | 41-[In]-27 |
| snR24 | snR24 | 63-39 |
| 5S | 5S | 20-1 |

**Supplementary Figure 1.**
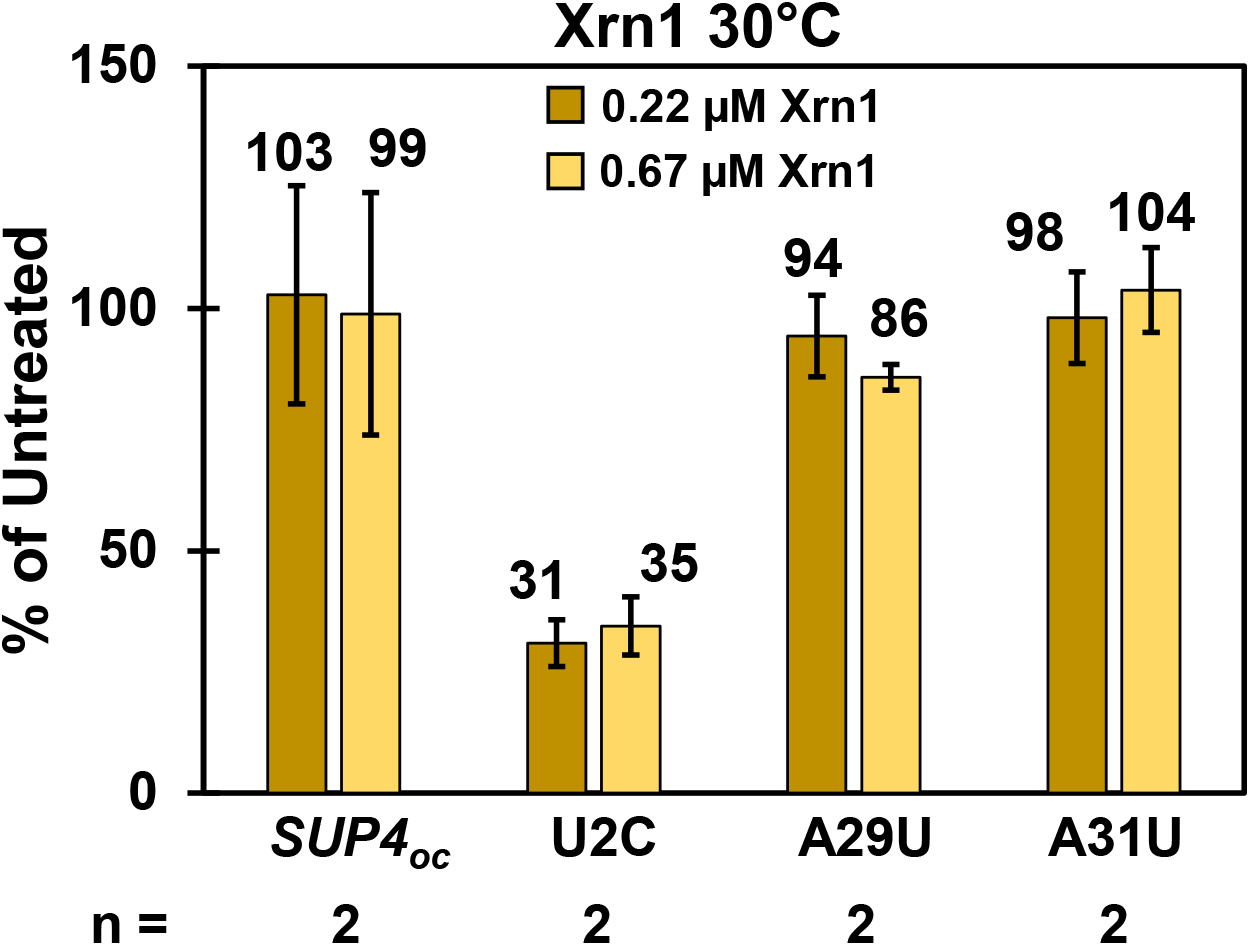
Anticodon stem variants are more resistant than an acceptor stem variant to *in vitro* degradation by Xrn1 at 30°C. **(A) Analysis of degradation of *SUP4_oc_* variants by purified Xrn1 at 37°C, experiment 2.** Degradation by purified Xrn1 was analyzed as described in Figure 2A using poison primer extension. The chart shows quantification of degradation by purified Xrn1 at 30°C, with data from two experiments conducted on different days

**Supplementary Figure 2.**
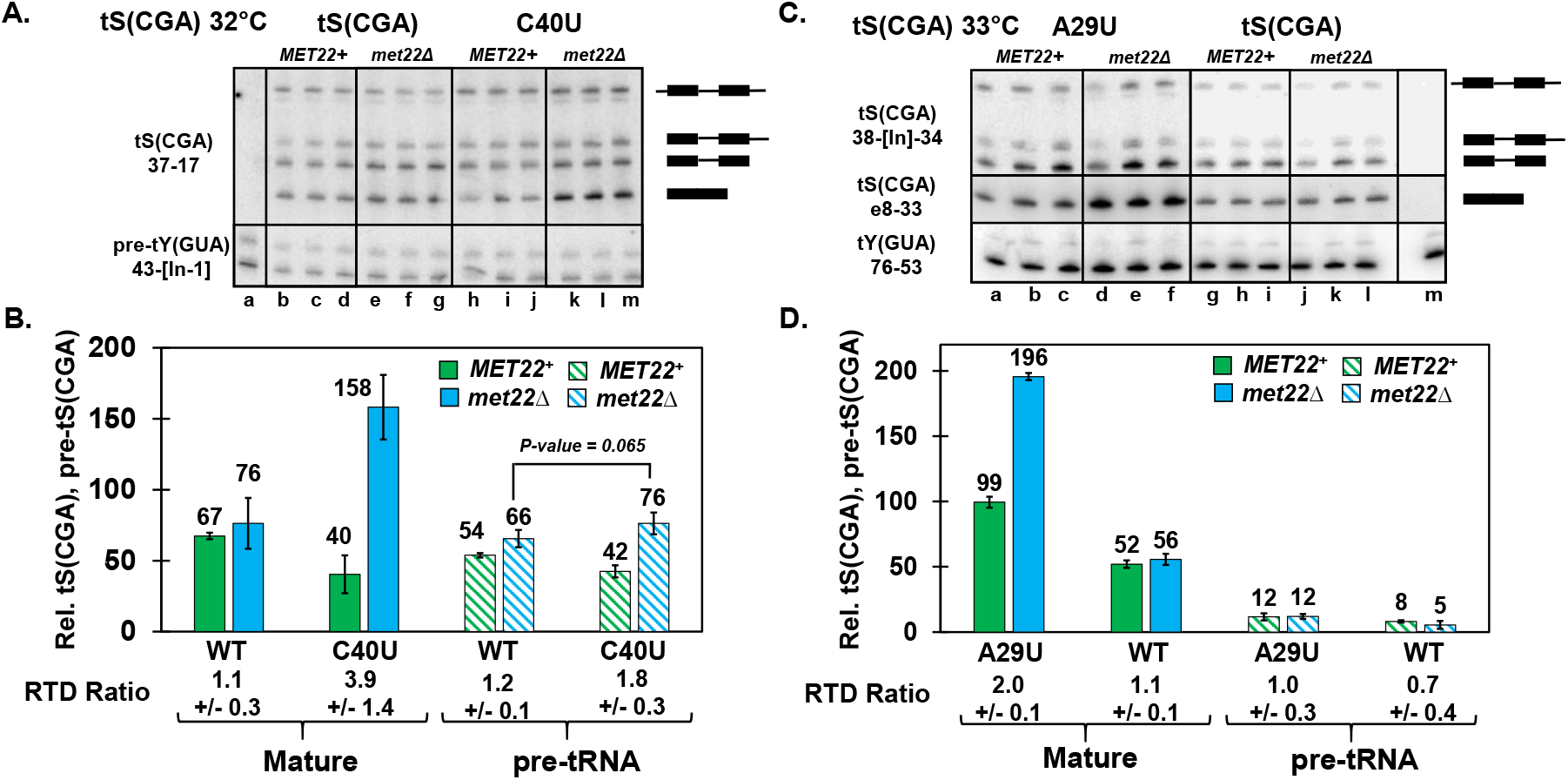
Analysis of RTD and pre-tRNA RTD in *tS(CGA)* variants. **(A) Northern analysis of *tS(CGA)* C_40_U tRNA levels in *MET22*^+^ and *met22Δ* strains grown at 32°C.** Strains with a *tS(CGA)Δ* mutation and a single integrated integrated *tS(CGA)* or *tS(CGA)* C_40_U variant as well as a *[tS(GCU/CGA) LEU2 CEN]* plasmid, were grown at 32°C, and bulk RNA was analyzed by Northern blot using probes with target regions as indicated. Lane a, strain with no *tS(CGA)*, used as a hybridization control. Pre-tRNA and mature tRNA species are indicated by cartoons. **(B) The *tS(CGA)* C_40_U variant is subject to RTD and modest pre-tRNA RTD.** Mature and pre-tRNA species were quantified relative to pre-tY to control for loading, and RTD and pre-tRNA RTD ratios are indicated at the bottom. The statistical significance of pre-tRNA accumulation was evaluated using a 1-tailed Student’s t-test assuming equal variance. **(C) Northern analysis of *tS(CGA)* A_29_U tRNA levels in *MET22*^+^ and *met22Δ* strains grown at 33°C.** Strains with a single integrated *tS(CGA)* or a *tS(CGA)* C_40_U variant were grown at 33°C, and bulk RNA was analyzed by Northern analysis as described in (A). Lane m, strain with no *tS(CGA)*, used as a hybridization control. **(D) The *tS(CGA)* A_29_U variant is subject to RTD but not pre-tRNA RTD.** Mature and pre-tRNA species were quantified relative to tY(GUA) to control for loading, and RTD and pre-tRNA RTD ratios are indicated at the bottom.

**Supplementary Figure 3.**
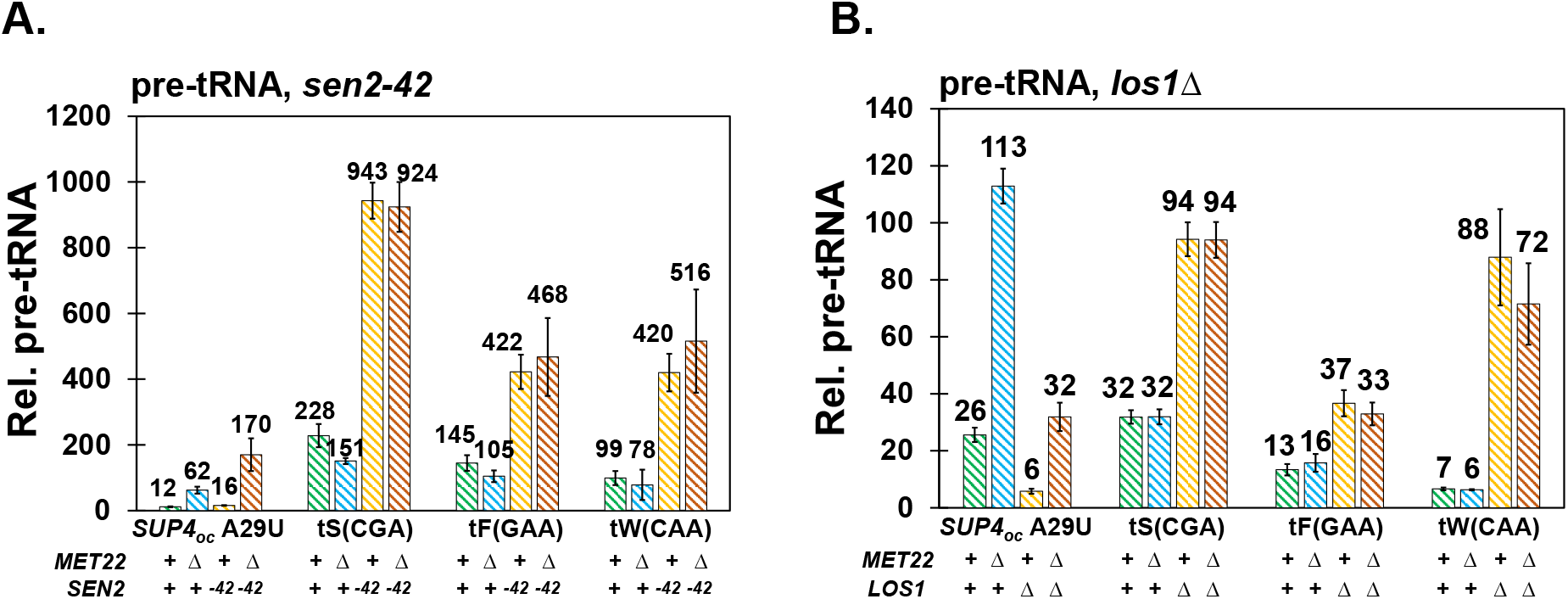
**(A) Quantification of pre-tRNA levels for the *SUP4_oc_ A_29_U* variant, compared to pre-tS(CGA), pre-tF(GAA), and pre-tW(CCA) levels, in *SEN2*^+^ and *sen2-42* backgrounds.** Bar graph of northern analysis in Figure 7A, right side. Pre-tRNA levels were quantified relative to hybridization of 5S to control for loading. **(B) Quantification of pre-tRNA levels for the *SUP4_oc_* A_29_U variant, compared to pre-tS(CGA), pre-tF(GAA), and pre-tW(CCA) levels, in *LOS1*^+^ and *los1Δ* backgrounds.**

